# Bioimage analysis for multiplexed FUCCI acquisitions powered by deep learning

**DOI:** 10.1101/2025.09.30.677886

**Authors:** J. Zimmermann, M. Pezzotti, E. Torchia, A. Enrico, M. Di Sante, F.S. Pasqualini

## Abstract

The FUCCI sensor fluorescently labels cell cycle phases, which is essential to assess normal and abnormal cell-cycle progression in physiological and pathological conditions of developing organisms. However, accurate cell-cycle decoding is challenging in the low signal-to-noise conditions typical of multiplexed live cell imaging. To address this challenge, we developed deep learning networks that integrate FUCCI signals with a cytoplasmic alpha-tubulin fluorescent reporter. Our approach outperforms existing methods for both segmenting and classifying FUCCI nuclei, even in low signal-to-noise conditions. The resulting high-accuracy segmentation enables robust automated tracking. We leverage this to introduce a dynamic time warping analysis that determines cell cycle pseudotime from incomplete tracks and can detect cell cycle arrest. We provide pre-trained networks for multichannel FUCCI analysis, offering a powerful tool for studies in cancer research, development, and mechanobiology.

## Introduction

The cell cycle is intrinsically linked to the cell fate and external cues acting on the cells. Stem cell fate is, for example, encoded in cell cycle progression patterns^1^ and the response of cancer cells to drugs^2^ or mechanical cues^3^ depends on the cell cycle. The visualization of the cell cycle progression in living cells is enabled by the widely adopted Fluorescent Ubiquitination-based Cell Cycle Indicator (FUCCI) (Fig. 1a)^4,5^. Multiplexing the FUCCI signal with other fluorescent reporters has recently been enabled^6^ and has unlocked the correlation of cell cycle information with cell-specific structural or functional information. However, a key challenge is robust bioimage analysis of FUCCI imaging data to obtain reliable quantitative information. Accurate cell-cycle decoding in multiplexed FUCCI is challenged by low signal-to-noise ratio (SNR) and spectral bleed-through. Specifically, the FUCCI sensor is expressed in the nucleus and emits in two colors with varying intensity depending on the cell cycle. Consequently, directly segmenting the FUCCI nuclei is challenging for many current bioimage analysis tools, which are typically trained on standard single-channel nuclear stains. Furthermore, multiplexed acquisitions in live imaging require low-power exposure to minimize photodamage to the cells, which results in a low signal-to-noise ratio (SNR)^7^, and is a challenge for pre-trained networks^8^.

**Figure 1:**
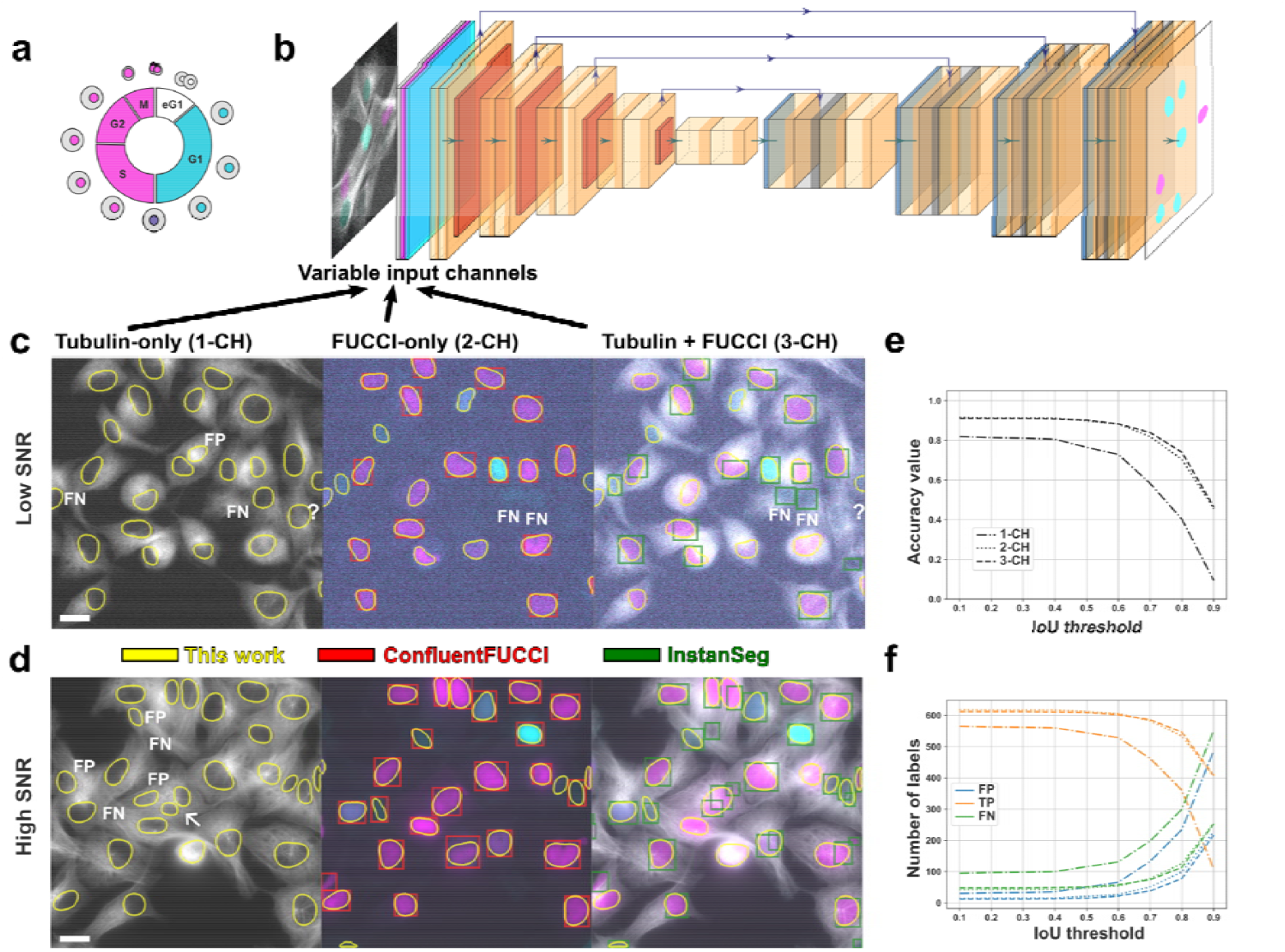
Segmentation workflow: **a** The FUCCI sensor allows to track the synthesis and degradation of fluorescently tagged proteins regulated by the cell cycle phases with distinct color combinations. In this work, the G1 phase is indicated by cyan, the S/G2/M phase by magenta, and the G1/S phase by both colors. **b** A U-Net-like segmentation network for nuclei with a different number of input channels is constructed depending on the available information from FUCCI imaging multiplexed with functional and structural sensors. **c** Segmentation results on a low SNR image of HT1080 cells are compared for the three trained networks with different input channels. The scale bar is 20 µm (bottom left). Results from two state-of-the-art methods are shown for comparison. Missing masks of our networks are denoted as FN (false negative), ambiguous masks by a question mark, and hallucinated masks by FP (false positive). **d** Similar to **c**, masks predicted on a high SNR image are shown and compared. One region, where the tubulin-only network predicted two masks instead of one, is highlighted by an arrow. **e** The accuracy over the intersection over union (IoU) threshold is shown for all three networks. **f** The number of false positive (FP), true positive (TP), and false negative (FN) labels are compared for the three networks.

To address this challenge, bioimage analysis pipelines have been devised that merge the two FUCCI channels to create a single-channel signal compatible with pre-trained deep learning segmentation networks. However, these solutions required retraining of the pre-trained networks^9,10^ to achieve the high segmentation accuracy required for reliable downstream analysis, such as cell tracking and cell cycle phase classification. Recently, channel-invariant networks have been suggested, which have not yet been explored for the segmentation of FUCCI data^11,12^. A further obstacle in the adoption of multi-channel or channel-invariant networks for FUCCI data is the lack of annotated imaging data.

We reasoned that training on data acquired in a heavily multiplexed acquisition scenario can help us solve these challenges. Since functional or structural sensors, such as the alpha-tubulin tag, are often expressed only in the cytoplasm we hypothesized that they could be used to aid in - or be sufficient for - conducting the nuclear segmentation task. Moreover, training a deep learning network on a multiplexed dataset with diverse SNR ratios should yield generalization and wide applicability on other low-SNR data.

To test this idea, we utilized a custom-trained convolutional neural network (CNN) based on StarDist^13^, which can accommodate up to three input channels. The CNN was trained on a diverse dataset of a human cancer cell line (HT-1080) and a keratinocyte cell line (HaCaT) imaged with various magnifications and SNRs. We tested the trained CNNs on challenging, low-SNR datasets and FUCCI data of other cell types from the literature. We observed that the segmentation task was performed well even when only the cytoplasmic signal was provided. We outlined how channel-invariant networks can be leveraged for multiplexed FUCCI data. We demonstrated that the high accuracy enables robust tracking of individual cells through the entire cell cycle. Furthermore, we introduced a post-processing algorithm to compare cells to the expected cell cycle progression. This approach enables the determination of the cell cycle percentage as a pseudotime, suggesting whether the cell follows the standard cell cycle without the need to acquire the entire cell cycle. The complete pipeline and training data are made openly available to provide researchers with a robust tool for their own studies.

## Results

### Deep learning for nuclear segmentation

We prepared a diverse dataset comprising fluorescent imaging data of HaCaT and HT1080 cells tagged with a custom FUCCI sensor^6^ (Fig. 1a), alpha-tubulin, and actin reporters. One channel of the FUCCI sensor has spectral overlap with the structural reporters (alpha-tubulin in case of the HaCaT cells and actin in case of the HT1080 cells). The cell seeding varied from individual cells to confluent layers (see also Fig. 4). Different magnifications (20x, 40x, 100x) and exposure conditions were used so that the dataset comprises low SNR acquisitions typical for live cell imaging as well as high contrast, high SNR acquisitions (Fig. 1c,d). The nuclei were segmented using a human-in-the-loop approach to prepare ground-truth data. Eventually, the dataset comprised about 5000 annotated nucleus instances from time lapses of more than 15 different samples.

To leverage the multiplexed acquisition, we trained a custom network with multiple input channel configurations (Fig. 1b). This choice was motivated by the observation that the alpha-tubulin signal contains information about the nuclear position. Hence, we used (i) only the alpha-tubulin channel (1-CH configuration), (ii) the two FUCCI channels (2-CH configuration), and (iii) the two FUCCI channels and the alpha-tubulin channel (3-CH configuration). We chose the StarDist architecture, which has been shown to segment nuclei highly accurately and can be straightforwardly modified due to its high-quality codebase^13^. We chose the accuracy at an IoU of 0.5 as a metric to evaluate the segmentation performance (more details in the Methods section)^13,14^.

On the validation data, we observed high accuracy with the custom-trained networks (see Fig. 1e and Table 1). The 2-CH and 3-CH networks achieved a similar accuracy of approximately 0.9. However, upon closer inspection, the 3-CH network detected nuclei that were missed by the 2-CH network, although it also missed some others that the 2-CH network found. In one dataset, we found that the 2-CH network misclassified bleedthrough artifacts as nuclei (see Video 1), whereas the 3-CH network segmented the images correctly. The one-channel network had an accuracy of 0.77. Upon visual inspection, we found that it accurately finds nuclei in tubulin structures that clearly delineate the nucleus, but can be misled when a hole in the tubulin network resembles a nucleus (see Fig. 1c). We also observed that occasionally a high tubulin signal is misidentified as a nucleus, as the tubulin signal increases during mitosis when cells round up.

**Table 1:**
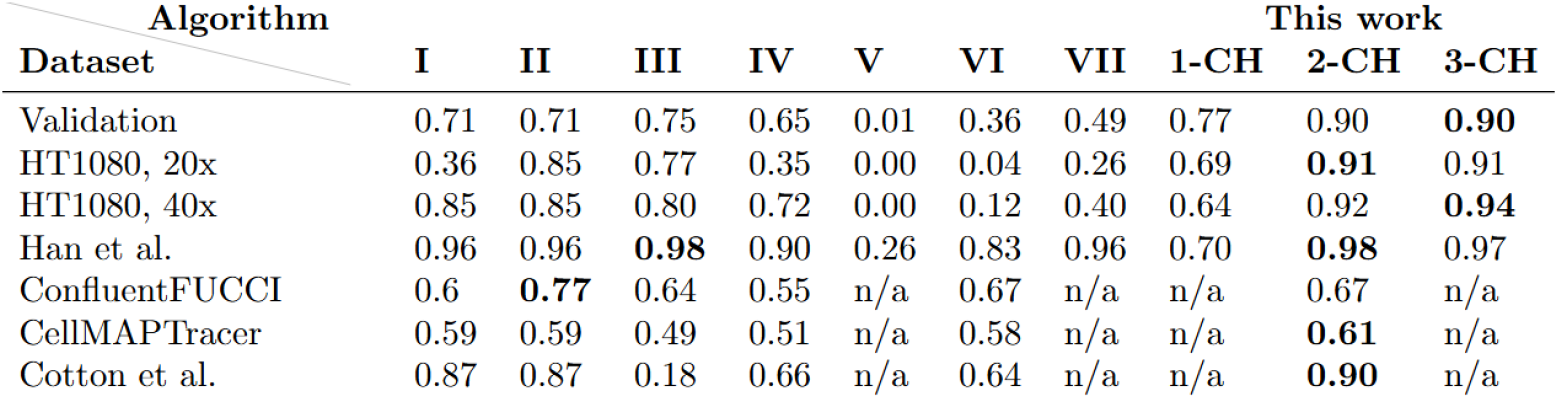
Segmentation accuracy at intersection over union of 0.5 on validation and literature datasets (more info in SI). Algorithms: **I** - DAPI-equivalent approach, no pre-processing, pre-trained StarDist segmentation, **II** - DAPI-equivalent, post-processing, pre-trained StarDist, **III** - DAPI-equivalent, post-processing, pre-trained generalist Cyto3 model, **IV**: pre-trained ConfluentFUCCI, pre-trained channel-invariant InstanSeg with tubulin-only input (**V**), **VI** - InstanSeg. pre-trained, 2-CH, **VII** - InstanSeg, pre-trained, 3-CH. Our models are denoted by **1-CH** (tubulin-only network), **2-CH** (FUCCI channels), and **3-CH** (tubulin + FUCCI channels).

An analysis of error types revealed that false positive (FP) segmentations, or erroneous predictions, occurred more often with the 1-CH network than with the other networks with a ratio of 8.6% (computed as the number of FP labels divided by the number of predicted masks). The ratio of FP predictions of the 2-CH and 3-CH networks was significantly lower with 3.3% and 2.4%, respectively. All networks had more false negative (FN) than FP segmentation masks, i.e., were rather missing than hallucinating nuclei (Fig. 1f). We also tested whether swapping the two FUCCI input channels impacted the segmentation accuracy. The accuracy dropped slightly by 0.02 due to a 3% higher rate of missed (FN) masks, indicating that channel order is not critical for the segmentation task and that the network does not heavily rely on morphological or intensity features specific to one FUCCI channel.

To benchmark our approach, we compared its segmentation accuracy against current state-of-the-art pre-trained solutions for FUCCI data and found that it matched or slightly outperformed them in high SNR and outperformed them in the low SNR regime (Table 1). We benchmarked our method against three approaches. The first combines the FUCCI channels into a single DAPI-equivalent channel^9^, which is then segmented by pre-trained StarDist or CellPose networks. The second consisted in the ConfluentFUCCI workflow, which treats the two channels separately and merges the segmentation masks in a post-processing step^10^. The third approach was the novel, pre-trained, channel-invariant InstanSeg network^11^, which directly segments multi-channel images without requiring pre- or post-processing. The pre-trained ConfluentFUCCI and InstanSeg networks performed well on high-contrast, high SNR images (Fig. 1c, d). Both approaches fail to segment masks with low FUCCI intensity in images with a noise level comparable to the FUCCI intensity. The DAPI-equivalent approaches were similarly unable to deliver accurate segmentations on our low signal-to-noise ratio (SNR) data. We tested the consistency of the trained network on a dataset of HT1080 cells that was acquired separately and was not part of the training and validation dataset (Table 1). Further, we investigated how performance depends on the imaging magnification. The pre-trained networks performed as expected on very noisy (20x magnification) and moderately noisy (40x magnification) data, outperforming all other benchmark methods.

To assess the generalizability of the network, we tested it on published FUCCI datasets. We annotated selected slices of the shared data to calculate the segmentation accuracy. On a high-quality dataset using the original FUCCI sensor in HaCaT cells^15^, we obtained a near-perfect segmentation result with the 2-CH and 3-CH network (accuracy of 0.97 and 0.98, respectively). However, all methods that considered the FUCCI signal performed only slightly worse on this high-quality data. This dataset^1^ included a YFP-SMAD2 reporter, which in the absence of the TGF-beta ligand is expressed in the cytoplasm^16^ and could therefore be used to segment the nucleus. Hence, we tested whether the 1-CH network trained on tubulin, which is also a cytoplasmic reporter, could segment the SMAD signal without modification. The accuracy was 0.7, slightly lower than the accuracy on our validation dataset but still outperforming the pre-trained InstanSeg network. The accuracy loss was mostly attributed to the low SMAD intensity in some cells (see Fig. S2b).

On other datasets containing only the FUCCI signal, we used our 2-CH network. We tested the data using the PIP-FUCCI sensor in RPE1-hTert cells^17^. On this dataset, the accuracy was significantly lower than on other datasets. The reason for this is that the signal from the PIP-FUCCI sensor becomes dark between the G1 and S phases, while the ground truth masks contain all nuclei identified by a separate reporter. Consequently, on a dataset using the PIP-H2A sensor developed to overcome this limitation^9^, our network again achieved high accuracy. In contrast to our validation dataset, the DAPI-equivalent approach also yielded promising results (accuracy of 0.87) on this dataset, comparable to those of our custom network (accuracy of 0.90, Table 1). These results suggest that the pre-trained networks can be used on pre-processed FUCCI data without a significant loss of accuracy. Our approach instead skips the pre-processing of the data.

Finally, to show that the success of our approach is not solely attributable to the StarDist architecture, we trained InstanSeg^18^ from scratch. The custom-trained InstanSeg networks reached approximately the same accuracy as our StarDist models. Furthermore, we utilized the 3-CH configuration to train the channel-invariant InstanSeg network^11^. This network performed similarly to the custom-trained, channel-specific networks when using the three channels, but it underperformed them when using less channels. With only one channel, the accuracy was almost zero. Likewise, we fine-tuned the recently proposed, channel-invariant transformer-based Cellpose-SAM network^12^. The custom-trained Cellpose-SAM network achieved a similar, but slightly worse, performance than our StarDist-based network when using all three channels. Using fewer channels reduced the accuracy, but Cellpose-SAM significantly outperformed the channel-invariant InstanSeg. However, we found that the inference time was at least ten times slower than InstanSeg and about five times slower than StarDist.

## Classification of cell cycle phases

Next, we explored whether deep learning can be leveraged to classify nuclei in single-frame acquisitions into the G1, G1/S, and S/G2/M phases, which are traditionally assigned based on the intensity of the FUCCI color combinations (Fig. 2a). The multiplexed acquisition introduces new challenges for the classification task because of spectral overlap. For example, the cyan intensity is elevated shortly before mitosis because the cells round up and the actin or tubulin signal bleeds into the nuclear cyan channel. This can lead to the misclassification of a cell in the S/G2/M phase as being in the G1/S phase.

**Figure 2:**
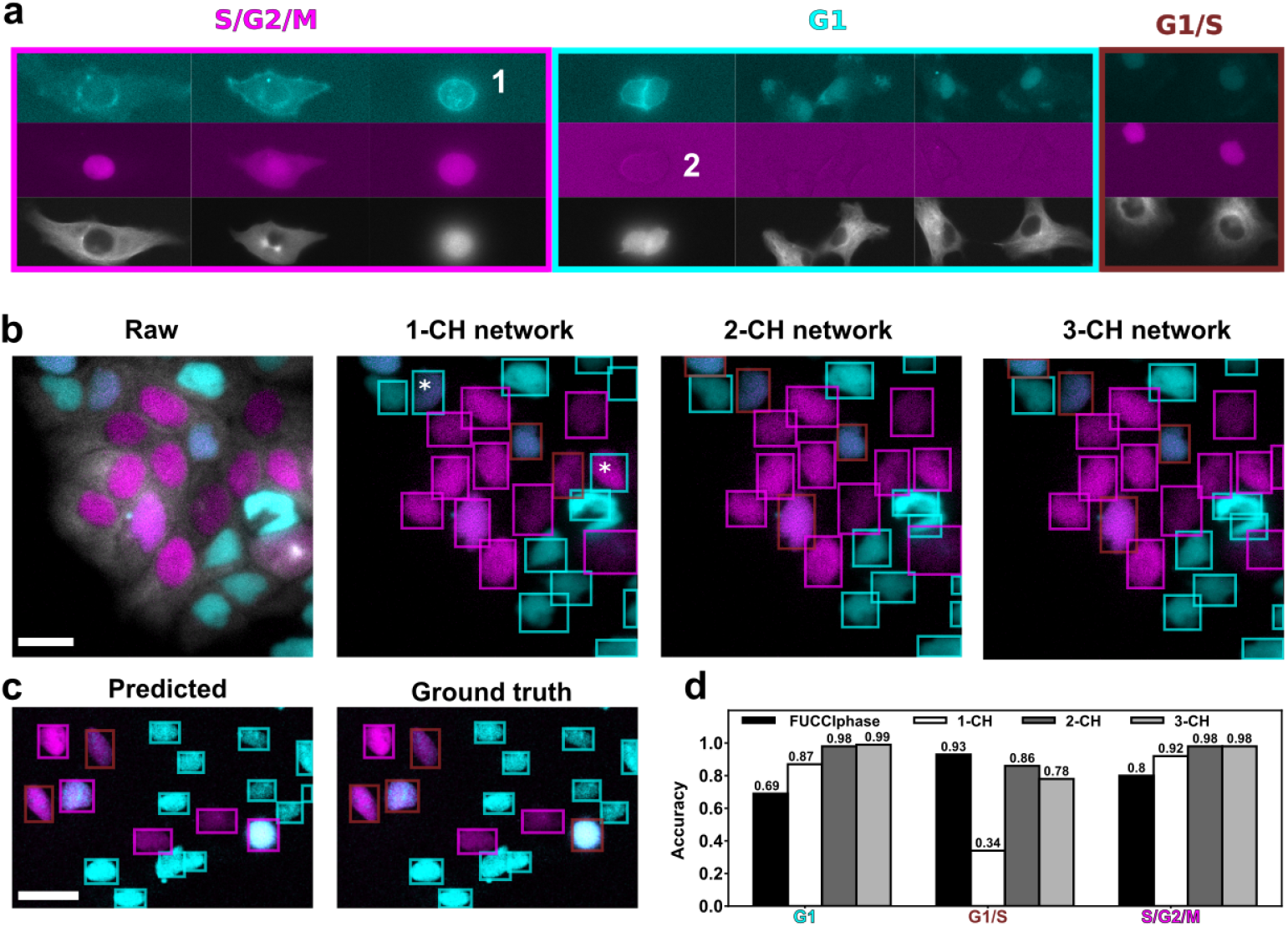
Classification workflow: **a** The classification into cell cycle phases is based on nuclear intensities of the two colors, here exemplified by HT1080 cells. The G1 phase is characterized by a high cyan intensity and no magenta intensity, the G1/S phase (labelled in brown) features both cyan and magenta intensity, and the S/G2/M phase is indicated by high magenta intensity and no cyan intensity. In multiplexed acquisitions, spectral overlap can lead to elevated intensities of colors that are not to be expected in the respective cell cycle phase. For example, there is a high signal in the cyan channel during mitosis (1), and autofluorescence is visible in the magenta channel in G1 phase (2). **b** Exemplary performance of the custom-trained classification network on HaCaT cells. Asterisks highlight misclassifications. Note that the difference between the 2-CH and 3-CH networks is only the segmentation. Bounding boxes were used for better visibility. **c** Performance of the classifier on the PIP-FUCCI sensor on literature data^17^. Here, the phases have been relabelled: cyan stands for the G1 phase, magenta for the S phase, and brown for the G2/M phase. **d** Comparison of the classification accuracy of the intensity-based classifier (FUCCIphase) and the 1-CH, 2-CH, and 3-CH networks on segmentation masks matched with the ground truth. All scale bars are 20 µm.

To train the network for the classification task, StarDist requires cell cycle phase labels for each segmentation mask. The classification network returns pixel-wise class probabilities, which are then aggregated for each segmentation mask. The loss function accounts for the class probability maps and differs from the loss function of the segmentation network ^19^. We compared the accuracy of the classification network to that of the segmentation-only network, both trained on the same data, to determine if the change in loss function affected its segmentation capabilities. The classification networks had a marginally lower accuracy than the segmentation network on the validation dataset, indicating that the added classification task did not significantly impair segmentation quality.

We assessed the network’s performance by calculating the class-wise accuracy and precision. We computed the precision and accuracy for each combination of predicted and ground-truth classes, yielding a matrix (Table 2). Here, precision indicates how often a mask was correctly predicted to be in the correct class, while accuracy also includes segmentation accuracy because it accounts for both correct and incorrect labels, including those resulting from erroneous predictions (FN). We expected high precision and accuracy values along the diagonal of the matrix and larger off-diagonal entries only when cell cycle phases are mistaken for each other. Our analysis showed that the FUCCI-only 2-CH network performs slightly worse at identifying G1/S nuclei, misclassifying them as G1 nuclei. Otherwise, it matches the 3-CH network in correctly identifying the phase of correctly detected nuclei. The 1-CH network, which relies exclusively on the tubulin channel, performs significantly worse but still identifies S/G2/M nuclei with reasonable success (see Fig. 2b), which could hint at S/G2/M-specific features contained in the tubulin network. G1 and G1/S nuclei were poorly detected and confused with each other. The low accuracy, which includes both the segmentation and classification performance, is as low as 0.26 for G1/S nuclei. An example showing the performance of the three networks on time-lapse data can be found in Video 2.

**Table 2:** Precision and accuracy of classifiers based on the tubulin-only (1-CH), regular FUCCI-only (first channel cyan, second magenta, 2-CH), swapped FUCCI-only (first channel magenta, second cyan, 2-CH), and three-channel network (cyan, magenta, tubulin, 3-CH). The swapped FUCCI configuration was used to check if the classification network focuses on the color combination.

| Precision |  |  |  |  |  |  |  |  |  |  |  |  |  |
| --- | --- | --- | --- | --- | --- | --- | --- | --- | --- | --- | --- | --- | --- |
| Prediction \ GT |  | Tubulin-only |  |  | Regular FUCCI |  |  | Swapped FUCCI |  |  | All channels |  |  |
|  |  | G1 | G1/S | S/G2/M | G1 | G1/S | S/G2/M | G1 | G1/S | S/G2/M | G1 | G1/S | S/G2/M |
| G1 |  | 0.70 | 0.34 | 0.08 | 0.92 | 0.08 | 0.00 | 0.02 | 0.05 | 0.88 | 0.92 | 0.03 | 0.00 |
| G1/S |  | 0.10 | 0.63 | 0.05 | 0.01 | 0.88 | 0.03 | 0.02 | 0.89 | 0.03 | 0.03 | 0.94 | 0.03 |
| S/G2/M |  | 0.06 | 0.00 | 0.84 | 0.01 | 0.04 | 0.96 | 0.91 | 0.06 | 0.04 | 0.01 | 0.03 | 0.95 |

| Accuracy |  |  |  |  |  |  |  |  |  |  |  |  |  |
| --- | --- | --- | --- | --- | --- | --- | --- | --- | --- | --- | --- | --- | --- |
| Prediction \ GT |  | Tubulin-only |  |  | Regular FUCCI |  |  | Swapped FUCCI |  |  | All channels |  |  |
|  |  | G1 | G1/S | S/G2/M | G1 | G1/S | S/G2/M | G1 | G1/S | S/G2/M | G1 | G1/S | S/G2/M |
| G1 |  | 0.49 | 0.03 | 0.03 | 0.79 | 0.01 | 0.00 | 0.00 | 0.00 | 0.73 | 0.82 | 0.00 | 0.00 |
| G1/S |  | 0.09 | 0.26 | 0.04 | 0.01 | 0.74 | 0.02 | 0.01 | 0.70 | 0.03 | 0.02 | 0.72 | 0.02 |
| S/G2/M |  | 0.03 | 0.00 | 0.74 | 0.00 | 0.01 | 0.92 | 0.80 | 0.01 | 0.03 | 0.00 | 0.00 | 0.90 |

To compare the pure classification performance against the conventional intensity-based approach, we considered only segmentation masks with an IoU greater than 0.5. The deep learning networks outperformed the conventional intensity-based classification (see Fig. 2d): The 2-CH- and 3-CH-networks reached almost perfect accuracy and even the 1-CH-network, which does not rely on FUCCI intensities, performed better than the intensity-based approach in identifying G1 and S/G2/M nuclei. In line with the previous result, G1/S nuclei were not well detected, and for this class, the intensity-based classification performed slightly better.

To determine whether the deep learning classifier bases its decision on nuclear intensities and the combination of the two FUCCI colors, we swapped the two FUCCI input channels. The results suggest that the network indeed mostly focuses on the nuclear intensity and color combination of the FUCCI channels (Table 2). For example, a nucleus is labeled as G1 when the nuclear intensity is high in the first channel and low in the second channel (for the other color combinations, see Fig. 1a and Fig. 2a). The accuracy dropped most significantly for nuclei in the S/G2/M phase, which indicates that the network also includes morphological features in its classification decision. This observation is likely explained by mitotic rounding and the related intensity changes due to spectral overlap. Given that the classification network seems to base its decision on the nuclear intensity and color combination of the FUCCI sensor, we hypothesized that the network could be repurposed for other two-channel FUCCI sensors with similar color combinations. One example is the PIP-FUCCI sensor^20^: G1 and S phases are marked by a single color, while the appearance of both colors marks the G2/M phase. To repurpose the classification network, the color combinations of our FUCCI sensor (Fig. 1a, Fig. 2a) need to be mapped to the PIP-FUCCI color combinations. The G1/S phase of our sensor, which exhibits a two-color appearance, corresponds to the G2/M phase of the PIP-FUCCI sensor. Likewise, the S/G2/M of our sensor corresponds to the S phase of the PIP-FUCCI sensor. The G1 phase is the same for both sensors. We visually compared the correspondingly relabelled predictions of our custom-trained network to ground-truth annotations on a PIP-FUCCI literature dataset^17^. We found that most predictions agree (Fig. 2c). This result confirms that the network focuses on the color combination of the nuclear intensities.

Finally, we compared the classification capabilities of the recently proposed transformer-based, channel-invariant Cellpose-SAM network^12^ to our CNN-based, channel-specific StarDist networks. We trained Cellpose-SAM on our multichannel dataset using all three channels during training and performed inference using the different color combinations, as in the validation of the StarDist-based network (Table 2). In other words, we trained one network and tested it with four different channel combinations without retraining. The channel-invariant classification network does not outperform the CNN-based classification network (Table S1), particularly when only the tubulin channel is used. When the Cellpose-SAM classifier was trained solely on the tubulin channel, the results improved but remained inferior to those of the CNN-based network, although it achieved a slightly higher accuracy for nuclei in the S/G2/M phase. Surprisingly, we found that the Cellpose-SAM classification network yields slightly better segmentations than the Cellpose-SAM segmentation network for all possible numbers of input channels. This may be explained by the fact that the segmentation and classification networks use different training parameters, which we did not further investigate or optimize in this context.

## Detection of cell cycle state by tracking

We tracked nuclei tagged with the FUCCI sensor and compared the obtained FUCCI intensity curves against a reference cell cycle (Fig. 3a,b) with the goal of inferring the cell cycle state. We previously demonstrated the reconstruction of the cell cycle percentage of HaCaT cells using dynamic time warping (DTW)^6^. Here, the cell cycle percentage serves as a pseudotime, revealing at which stage of the cell cycle an individual cell is. DTW distorts the time axis to minimize the Euclidean distance between the query (tracked FUCCI intensity) and the reference curves (see Fig. 3c,d). To compensate for intensity differences between individual nuclei, we processed the signals using smoothing and z-score normalization, and then performed the distance computation on the signal derivative.

**Figure 3:**
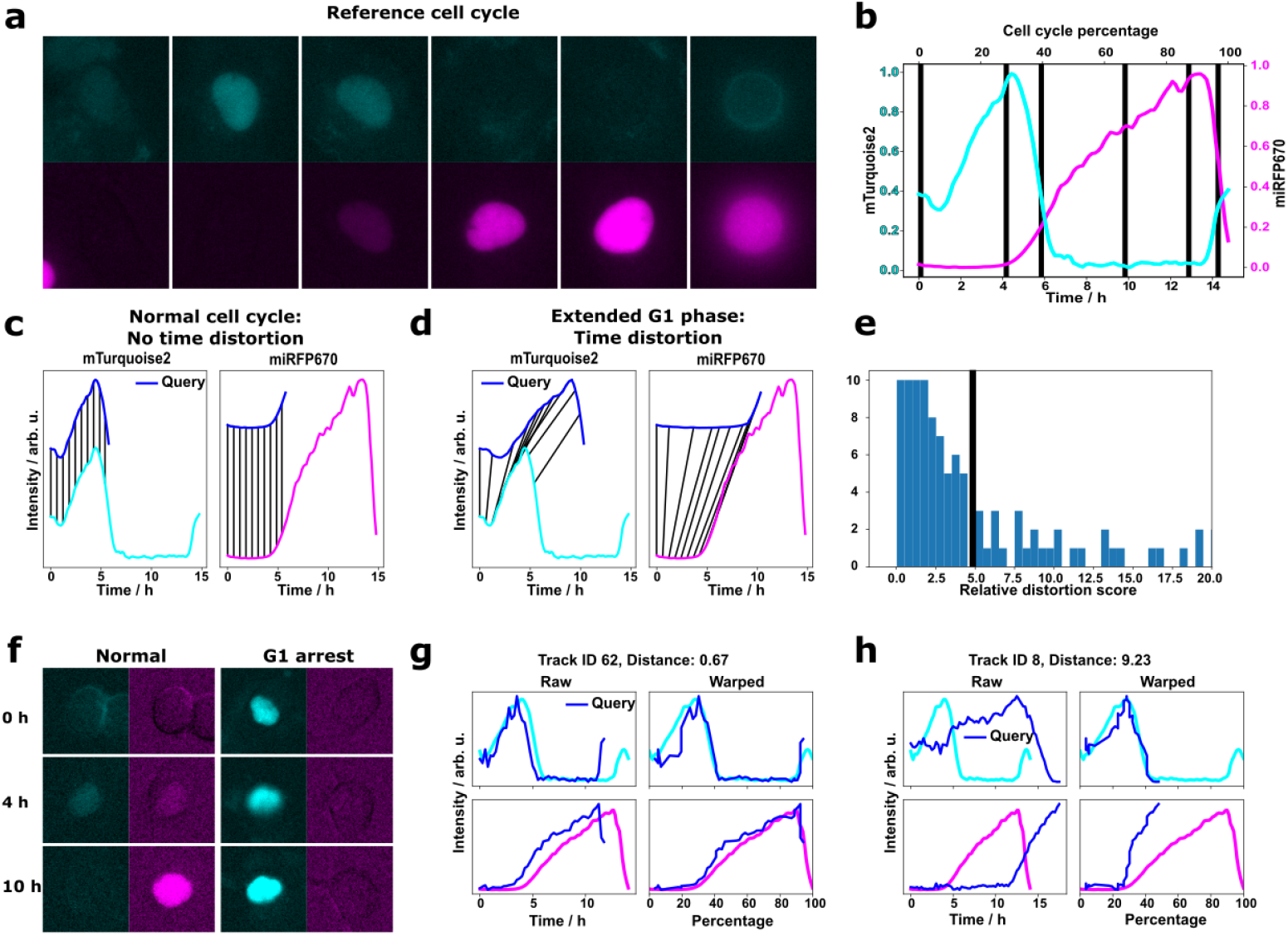
Tracking analysis for FUCCI data: **a** The FUCCI intensities change throughout the cell cycle. **b** They can be averaged for multiple cells to obtain a reference curve encoding both the absolute time of the cell cycle and the cell cycle percentage as a pseudotime. **c** Reconstruction of the cell cycle percentage is possible by querying the reference cell cycle using dynamic time warping (DTW). **d** If the query does not match well with the reference, the DTW algorithm distorts the signal along the time axis. **e** The relative time distortion value as a distance metric reflects the agreement with the reference cell cycle. By manually thresholding the histogram (black vertical line), we identified two sub-populations of HT1080 cells. **f** Cells with a high score were G1-arrested. A comparison of the warping process for a normal and a G1-arrested cell is shown in **g** and **h**. The relative time distortion value was used as the distance measure.

We leveraged the accuracy of the 3-channel segmentation network, which permitted us to automatically process low-SNR videos of HT1080 cells imaged at 20x magnification (see also Video Fig. 1). The few wrong segmentation masks did not necessitate manual corrections because they usually appeared in individual frames and were not linked by the tracking algorithm. The HT1080 cells could be distinguished based on their cell cycle state: in addition to cells following the normal cell cycle, some cells were arrested in the G1 phase (Fig. 3f). Warping the FUCCI intensity curves of G1-arrested cells on the reference curve by DTW requires a strong distortion of the signal along the time scale. This time distortion can be quantified by a recently introduced metric ^21^, which measures the stretching and compression of the intensity curves. We divided this metric by the number of frames to make it comparable between tracks of different length and refer to it as the relative time distortion value. By applying a threshold to this metric, we could distinguish between normally cycling and arrested cells based on the DTW alignment of their tracked FUCCI intensities (Fig. 3e, g, h). A relative time distortion value of 5 was manually defined to separate normal cells from G1-arrested cells. An example of a mixed population of cells following the normal cell cycle and G1-arrested cells, and the respective reconstructed cell cycle percentages is shown in Fig. 4a and Video Fig. 3.

**Figure 4:**
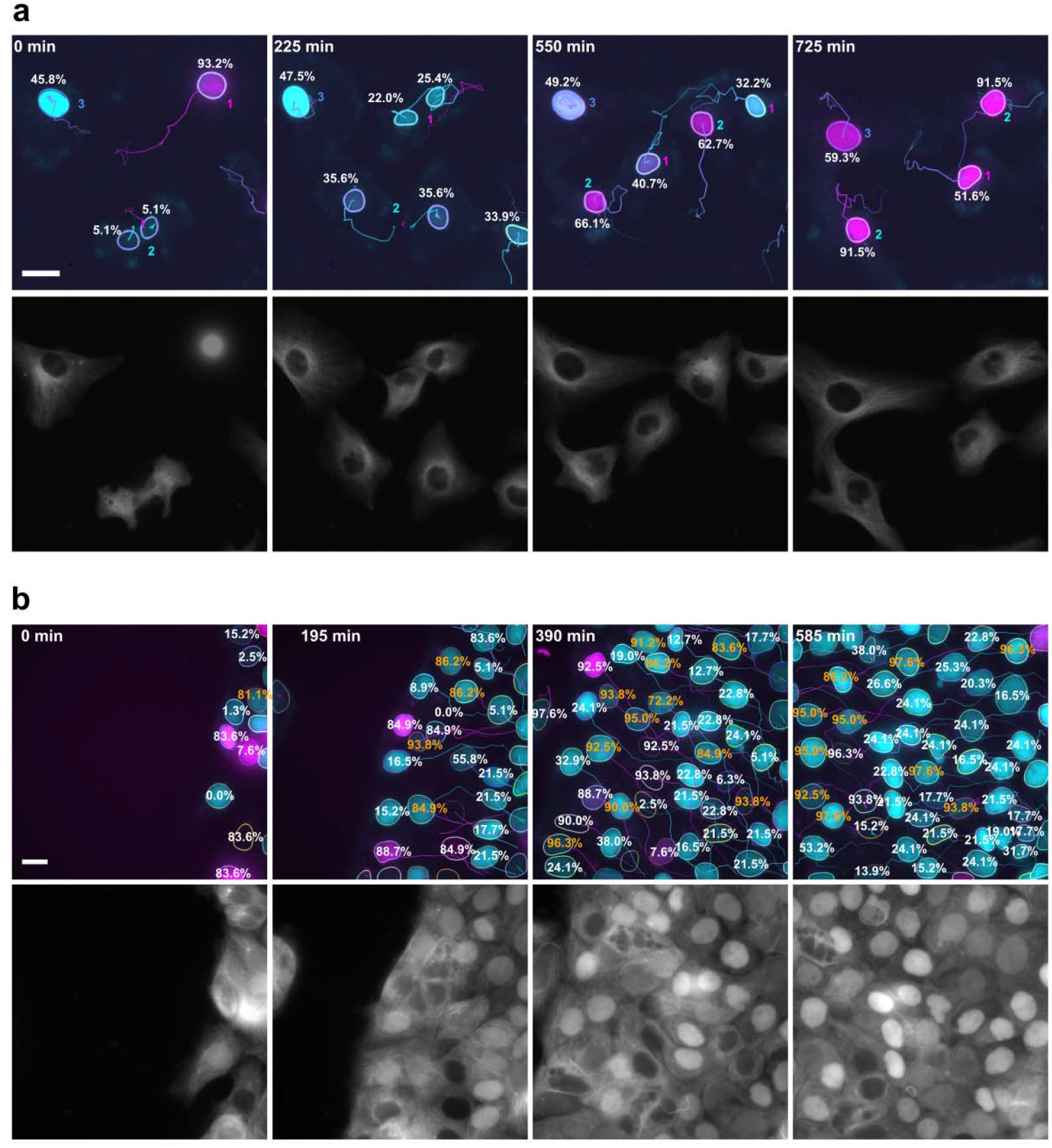
**a** Example of HT1080 cells annotated with cell cycle percentages. The numbers indicate the mother cell of the nuclei. Nucleus 3 is G1-arrested, while the other cells cycle normally. The tubulin channel is shown separately. **b** Example of HaCaT cells in a scratch assay experiment imaged with 100x magnification. Wrongly estimated cell cycle percentages. Note that compared to Video Fig. 4 some labels of cells at the image borders and short tracks have been removed for the sake of visibility. The tubulin channel clearly shows the spectral overlap with the cyan channel. All scale bars are 10 µm.

We tested the approach with confluent HaCaT cells in scratch assay, where we also expected extended G1 phases due to contact inhibition. Again, the approach worked out of the box, but revealed a limitation of the DTW approach stemming from its reliance on relative signal dynamics rather than absolute intensity. Because the DTW alignment is performed on normalized derivatives, the mapping has no information about the absolute FUCCI intensities. This can lead to failed cell cycle percentage reconstructions: In some cases, cells that were clearly in the G1 phase were mapped to a pseudotime at the end of the cell cycle (Fig. 4, Video Fig. 4). This issue primarily affected cells in the G1 phase that remained longer than expected in this phase. We rarely observed this issue in the HT1080 cell population. Despite the wrong cell cycle percentage reconstruction, these cells can be identified as not adhering to the standard cell cycle based on the DTW distortion, which is large for these cells.

## Discussion

Processing multichannel FUCCI data is difficult with conventional bioimage analysis tools because low-SNR fluorescence and spectral bleed-through degrade the nuclear signals and hamper downstream quantification. Deep learning offers a viable alternative, yet most pre-trained networks struggle with nuclear segmentation on multichannel low-SNR data, and channel-invariant approaches have only recently been introduced ^11,12^. Furthermore, classification networks must be trained for each specific task. We address these challenges by demonstrating how multichannel FUCCI data can be directly processed using deep learning, leveraging multiplexed acquisitions that comprise a structural sensor in addition to the FUCCI sensor^6^. We trained StarDist-based networks with a specific number of input channels to enable segmentation of nuclei throughout the entire cell cycle. Incorporating the cytoplasmic alpha-tubulin reporter in addition to the nuclear FUCCI signal was critical to segment nuclei with a dim FUCCI signal under low-SNR imaging conditions. The challenge of a dim FUCCI signal is also present in adaptations such as the PIP-FUCCI sensor^20^, where the intensity of both channels is dim between the G1 and S phases. Previously, this challenge has been addressed by introducing a specific nuclear stain^17^, i.e., adding another fluorescent nuclear sensor, which limits the possibility to multiplex the FUCCI signal with functional or structural reporters. An alternative FUCCI construct has also been suggested^9^, but it does not permit cell cycle phase determination from single snapshots. Our approach of incorporating a structural or functional marker into the segmentation network addresses this challenge while also providing access to additional phenotypic information. Remarkably, the nuclear segmentation task performed better than conventional approaches on the validation dataset even when using only the cytosolic alpha-tubulin reporter. It also showed promising results on test data sets, including one using the cytoplasmic SMAD reporter instead of the tubulin sensor. These results suggest that identifying nuclei from a cytoplasmic reporter, such as alpha-tubulin, could be leveraged to reduce phototoxicity by imaging the two FUCCI channels less frequently than the tubulin channel, without compromising trackability.

Our dataset comprised two cell lines imaged under various conditions. To assess the generalization of our 2-CH FUCCI segmentation network, we segmented literature data from other cell lines with various SNR values expressing the conventional FUCCI sensor^5^, the PIP-FUCCI^20^, and the PIP-H2A construct^9^ (Table 1). Our network performs as well or better than the benchmark methods, which highlights the general applicability of our network on two-channel FUCCI data. If necessary, the data can then be manually adjusted to expand our openly shared dataset and retrain the network using the extended dataset. To our knowledge, there exist no other publicly available networks pre-trained for multichannel FUCCI segmentation. Channel-invariant networks are promising when channel availability varies. In this work, we demonstrated their potential by training the recently released Cellpose-SAM^12^ and InstanSeg^11^ networks. Cellpose-SAM was less sensitive to using fewer channels in inference than in training (e.g, using one or two channels in inference, but three in training). Nevertheless, it requires significantly more training resources, performs slower inference (approximately five times slower than StarDist and approximately ten times slower than InstanSeg), and does not significantly outperform the channel-specific CNN-based networks with the available dataset. Larger and more diverse datasets may change this balance.

The identification of the cell cycle phase from FUCCI data is usually performed by analyzing FUCCI intensity per segmentation mask, either by thresholding^22^ or training a separate classifier ^23^. We investigated semantic segmentation, in which the nucleus is segmented and classified by the same network^19^ (Fig. 2). Our results show that this approach reliably performs the classification task and outperforms conventional intensity-based methods. We further found that the network primarily relies on FUCCI intensity information, which enabled us to classify data using the PIP-FUCCI sensor that had not previously been known to the network. Instead, the classification using only the structural alpha-tubulin reporter turned out to be unreliable. Unlike in the segmentation case, it gives limited access to the cell cycle state. It could become a subject of future research to investigate whether other reporters are more suitable for this task. Similar to the segmentation task, channel-invariant networks like Cellpose-SAM may provide an alternative to current CNN-based networks in future research; however, they do not outperform them at the current stage and with the currently available data.

The cell cycle can vary significantly between cells. Hence, the FUCCI sensor has been used to understand the cell cycle-related aspects of self-renewal and differentiation of stem cells^1^, and the impact of drugs on cancer cells^2^. To date, such analyses have required tracks spanning the entire cell cycle. We propose an approach to estimate the cell cycle percentage as a pseudotime from FUCCI intensity sequences that do not necessarily span an entire cycle (Fig. 3). We leverage DTW, which, to our knowledge, has not previously been used on FUCCI data, but has been used for cell cycle phase identification from histone labels with a focus on mitotic phase identification^24^. Importantly, our approach also allows us to quantify the agreement with a reference cell cycle. Using this, we could detect G1-phase arrest in HT1080 cells by thresholding the relative time distortion value of the DTW alignment. This capability is relevant for cancer research, where it has been shown that proliferative cancer cells are less invasive^3^ and that cancer drugs affect cells differently depending on the cell cycle state^2^. The presented approach compares relative intensity values, which we addressed through normalization and differentiation. The resulting limitation is that the DTW mapping has no notion of absolute intensities, which can lead to failed reconstructions, as experienced with HaCaT cells that have an extended G1 phase. In future research, sequence models that operate directly on the imaging data could be considered^25^. Such models would require extensive training datasets, which our reference-curve approach avoids, as a reference can be obtained from a single cell track.

While we presented our workflow only for 2D data, the approach can be straightforwardly extended to 3D data, which StarDist natively supports^26^. The main challenge will be to provide a sufficient amount of annotated training data. Our 2D network could be useful for facilitating this process by providing initial segmentations to generate consensus 3D segmentation masks^27^. Furthermore, we did not focus on tracking accuracy in this work. However, providing robust segmentation for all nuclei, as our method does, is the essential first step for any tracking algorithm. In addition, providing multiple segmentations with our multichannel networks could benefit the tracking accuracy, as recently demonstrated^28^.

In conclusion, we expect our solution to benefit researchers studying the impact of the cell cycle using multiplexed FUCCI sensors in low-SNR conditions in fields such as stem cell research^1^, cancer drug discovery^2^, and mechanobiology^29^. Our multi-channel open-source network automatically segments and classifies nuclei with high accuracy throughout the entire cell cycle, which in turn enables robust automated tracking of the cell cycle phase. Our automated DTW-based approach identifies cell cycle events from partial tracks, unlocking a new level of analysis for FUCCI-based studies by linking cell cycle dynamics to other structural or functional reporters.

## Materials and Methods

### Cell lines

We used the epithelial HaCaT clonal cell line as described in our recent work^6^.

The cells were genome-edited to express the FUCCI cell cycle indicator (mTurquoise2 (CFP) and miRFP670 (iRFP) fluorophores), to tag alpha-tubulin with an EGFP fluorophore at the N terminus, and to tag actin with the RFP LifeAct sensor.

This cell line has a spectral overlap between the EGFP and CFP fluorophores, i.e., between alpha-tubulin and nuclei in G1 phase.

In addition, we used the immortalized human fibrosarcoma HT1080 cell line (Ibidi, #HT-1080-LifeAct-TagGFP2, catalog 40101), which expressed the same fluorophores (more details can be found elsewhere^30^). Here, actin was tagged with the EGFP fluorescence protein. In addition, endogenous α-tubulin (TUBA1B, NM_006082.3) was tagged with tagRFP using the Thermo Fisher Scientific TrueTag system. Thus, the spectral overlap in genome edited HT1080 is between nuclei in G1 phase and actin.

Both cell lines were maintained in Dulbecco’s Modified Eagle Medium / Ham’s F-12 Nutrient Mixture without phenol red (DMEM F-12, Gibco, catalog# 21041-025), supplemented with 10% heat-inactivated fetal bovine serum (FBS, Gibco, catalog# 10270-106) and 1% Penicillin–Streptomycin (Himedia, catalog# A001-100ML). Cells were cultured at 37 °C in a humidified 5% CO□ incubator and routinely split at 70% confluency. For experiment preparation, cells were detached using Trypsin/EDTA 0.25% (Thermo Fisher Scientific, catalog# 25200056), resuspended in complete DMEM F-12, and counted adequately according to the experimental needs.

### Fluorescence microscopy

The cells were imaged using a Crest V3 X-Light spinning disk confocal microscope (Nikon) equipped with a Celesta Light Engine source (TSX5030FV, Lumencore), and a Photometrics Kinetix Scientific CMOS camera as in our previous research ^6,31^.

HT1080 widefield imaging at 20× was performed using sequential 638, 546, 477, and 446 nm excitations with exposure times optimized per channel in the 300–20 ms range.

HaCaT cells were imaged as described in previous works^6,31^.

The emitted signal was filtered using single band FF01-484/561 (catalog #FL-412124, Semrock) for the mTurquoise2 signal, single band FF01-685/40-25 nm (catalog #FL-011482, Semrock) for the miRFP670 signal, single band FF01-595/31 (catalog #FL-004391, Semrock) for the RFP signal, and FF01-511/20-25(catalog # FL-004306, Semrock) for the EGFP signal.

Different objectives with different magnifications were used: a Nikon CFI Plan Apo 20X objective (air-immersion, N.A. 0.75, WD 1 mm, catalog# MRD00205, Nikon), Nikon CFI Plan Apo Lambda S 25XC Sil objective (silicone oil immersion, N.A. 1.05, WD 0.55 mm, catalog# MRD73250, Nikon), a Nikon CFI Plan Apo Lambda S 40XC Sil objective (silicone oil-immersion, N.A. 1.25, WD 0.3 mm, catalog# MRD73400, Nikon), and a Nikon CFI SR HP Plan Apo Lambda S 100XC Sil objective (silicone oil-immersion, N.A. 1.35, WD 0.31 – 0.28 mm, catalog# MRD73950, Nikon). The immersion oil was Nikon silicon immersion oil 30cc (catalog #MXA22179, Nikon).

The pixel size of the camera is 6.5 µm and at max a 2700×2700 pixels field-of-view was recorded. The smallest pixel size in the data set was 335 nm with 20x magnification.

All images were acquired in 12-bit mode by Nikon NIS elements.

If applicable, the images were flat-field corrected using BaSiCpy ^32^.

### Data annotation

For simplicity, we will refer to the G1-phase channel as cyan channel and to the S/G2/M-channel as the magenta channel. The dataset was annotated in a human-in-the-loop workflow. Initial segmentation masks were generated by the DAPI-equivalent approach, which involves pre-processing of the cyan and magenta channel by denoising using a median filter, background subtraction using a top-hat filter implemented in pyclesperanto based on Clij ^33^, and a maximum projection to obtain one DAPI-equivalent nuclear channel. The DAPI-equivalent channel was segmented using a pre-trained StarDist model ^13^, i.e., required no in-house training. The masks were then displayed with the cyan, magenta and alpha-tubulin channel and manually curated in Napari ^34^.

Multiple subsequent frames were annotated together to avoid wrong annotations and to take the information about the cell cycle into account. This was necessary because the sensor has a very low intensity in the first frames before and after mitosis. In this case, the tubulin channel yielded information about the nuclear location (Fig. 1). Later, intermediate versions of the custom-trained networks were used for the human-in-the-loop approach. This reduces the annotation time for researchers using the network on their own cells.

The cell cycle phases were prepared using the fucciphase package ^6^, saved in JSON format, and manually corrected. A custom Python script was developed to facilitate this task.

Finally, the imaging data was scaled to the smallest pixel size (if needed) and tiled into crops of at least 256×256 pixels. As a result our dataset is trained on images with a pixel size according to 20x magnification, i.e. about 335 nm. We removed crops that did not have at least 4 labels inside the field of view (labels touching the boundaries were not counted).

For the test datasets (HT1080 20x and 40x), we used intermediate versions of the custom-trained networks and manually curated the labels. The datasets contained videos of 9 frames, only the middle frame was curated and the other frames were used to check the correctness of the segmentation as previously described.

### Training and validating the deep learning segmentation and classification

The dataset was randomly split into 85% used for the training and 15% used for the validation. We trained StarDist (version 0.9.1) using 200 steps per epoch and 1000 epochs. Random flips, rotations, intensity changes and Gaussian noise were used for data augmentation. For the classification task, we did not perform any additional augmentation. The typical StarDist configuration was used ^13,19^: the Adam optimizer was used with an initial learning rate of 3e-4 and a learning rate scheduler that halved the learning rate when the loss did not change for 80 epochs. The final model is the model with the smallest validation loss.

InstanSeg was trained with the settings as specified by its developers using random flips and rotations as data augmentations following the example online (https://github.com/instanseg/instanseg). Cellpose-SAM was trained following the example script provided online using 200 epochs (https://github.com/MouseLand/cellpose).

For comparison, we used the previously established DAPI-equivalent approach with and without pre-processing and with StarDist (“2D_versatile_fluo” model) or Cellpose (v 3.1.1.2, “cyto3” model^8^) as backend, respectively. The ConfluentFUCCI approach segments both FUCCI channels separately using a custom-trained Cellpose network (also used with Cellpose v 3.1.1.2). ConfluentFUCCI in its original formulation uses tracking data to identify correct segmentation masks. As we worked on a single-frame solution, we merged the segmentation masks using the Clij library^33^ in Python through the pyclesperanto_prototype interface.

To quantify the segmentation accuracy, we computed the mean accuracy at a given intersection over union (IoU), also known as average precision, as TP / (TP + FN + FP), where TP are the true positive labels, FN the false negatives and FP the false positives ^13^. Unless otherwise stated, we evaluated the accuracy at an IoU of 0.5. For the classification, we additionally considered the precision defined as TP / (TP + FP). The intensity-based classification was conducted on the segmentation masks obtained with the 2-CH network using the fucciphase package and setting a threshold of 0.1 with respect to the maximum intensity.

We used different datasets that are described in greater detail in the supplementary material.

The training and validation was performed on a server with two NVIDIA RTX A6000 GPUs, 512GB RAM, and two AMD EPYC 7763 64-core processors. The networks were used on a workstation with an NVIDIA GeForce RTX 4070 GPU, 128 GB RAM, and an AMD Ryzen 9 7900X3D 12-core processor and on a laptop with a NVIDIA GeForce RTX 3050, 32GB RAM, and a Intel i7-12700H processor.

### Cell tracking and postprocessing

The image data must have a constant background intensity in all frames. Otherwise, the extracted FUCCI intensities do not reliably reflect the cell cycle. Thus, we either corrected the background intensity by time-lapse processing in BaSicPy ^32^ or performed a frame-wise percentile normalization when no flatfield correction was applied. The necessity of the flatfield correction was decided based on line profiles of the background intensity. At low magnifications(20x, in some instances also at 40x), vignetting was observed and corrected through the flatfield correction.

The image data were augmented by an additional channel, which contained the segmentation masks. The augmented image data were loaded into Fiji ^35^ and processed using the TrackMate plugin ^36^. The label detector was used on the channel containing the segmentation masks. The standard LAP tracker was used with settings that were manually adjusted to enable a good linking quality.

The tracks were postprocessed by the TrackMate actions “*Close gaps in tracks by introducing new spots”*, which introduced new spots in tracks by interpolation, and *“Auto naming spots”*, which appends letters for each branch so that cell divisions can be detected from the spot name. The TrackMate tracking results were exported as XML files and further postprocessed with the fucciphase Python package. To match the tracked FUCCI intensities with the reference intensity curve, we used subsequence matching as implemented in DTAIdistance ^37^. This approach was already described for the case of HaCaT cells in our earlier work ^6^. In this work, we implemented the time distortion coefficient recently described ^21^ in fucciphase. Because we match subsequences of varying length, the time distortion coefficient was divided by the track length.

The reference curve of the FUCCI intensity for HT1080 cells was extracted from four tracks of HT1080 cells imaged with 40x magnification (Fig. 3a, b). The reference curve for HaCaT cells was taken from our previous work^6^, where it was extracted from 11 tracks.

## Acknowledgements

This work was supported by the European Research Council (ERC) Starting Grant No. 852560 to F.S.P. and by the Italian Ministry of Education, University, and Research (MIUR) (FARE2020, Grant No. R20ZE54CTK) to F.S.P.

We thank Haris Iqbal for sharing LaTex code for the visualisation of the U-Net architecture: https://github.com/HarisIqbal88/PlotNeuralNet/

## Data and code availability

The code and instructions to prepare and process the data can be found on GitHub: https://github.com/synthetic-physiology-lab/deepfucci

The fucciphase package is also available on GitHub: https://github.com/Synthetic-Physiology-Lab/fucciphase

The annotated datasets, pretrained models, and code repositories at the time of submission were deposited on Zenodo and are accessible under <u>this link</u> (the final repository will be published upon acceptance).

## Author contributions

J.Z. conceptualized the study together with F.S.P., wrote all software, annotated the data, and wrote the initial draft of the manuscript. M.P., E.T., A.E., M.D. collected the data. F.S.P. supervised the project and acquired funding. All authors have reviewed and edited the final version of the manuscript.

## Footnotes

1 https://github.com/Zi-Lab/eDetect/releases

