## Supplementary Information for "Bioimage analysis for multiplexed FUCCI acquisitions powered by deep learning"

#### HT1080 test dataset (own dataset)

The HT1080 cells were imaged at different levels of confluency with 20x magnification (746 nuclei) and 40x magnification (734 nuclei) (see Fig. S1a and S1b, respectively).

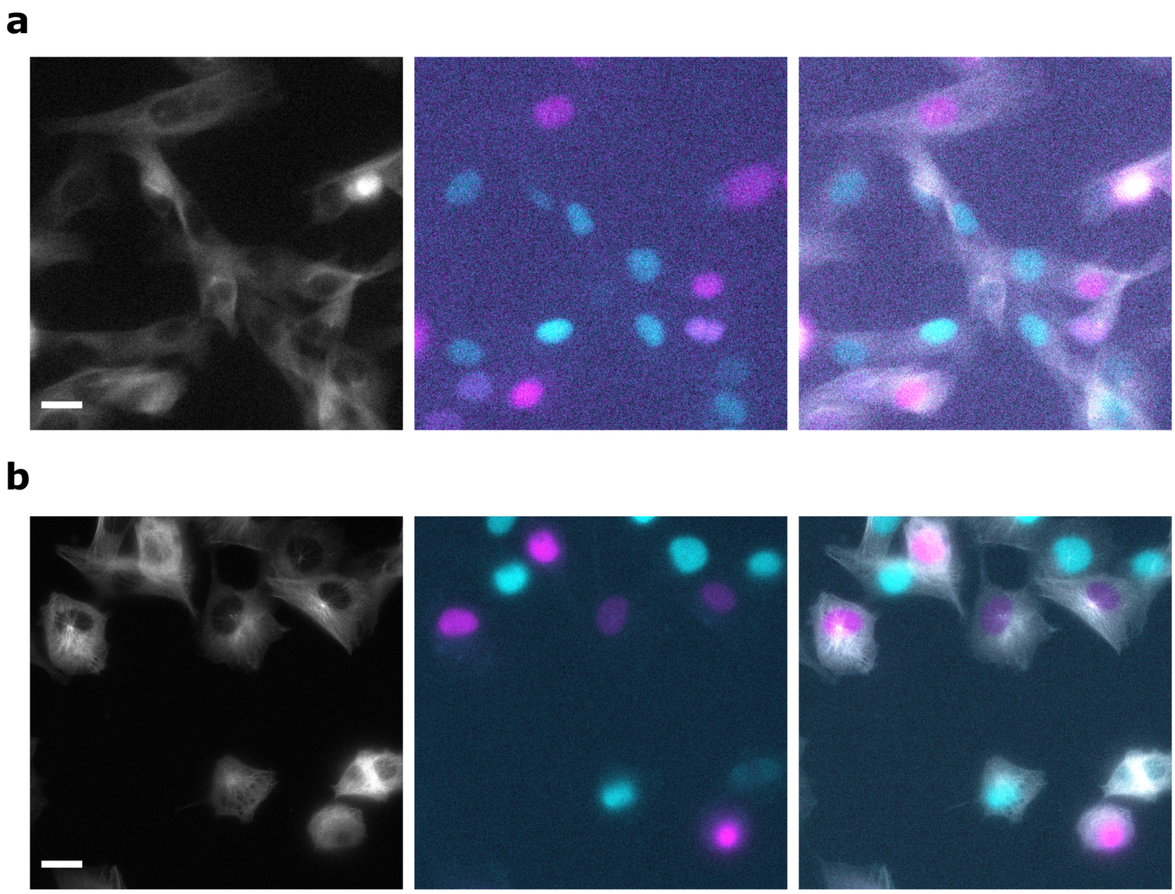


Figure S1: Exemplary HT1080 dataset used to test the segmentation network: Cells were imaged with 20x magnification (**a**) and 40x magnification (**b**). The scale bar is 20 µm.

### Han et al.[1]

HaCaT cells labelled with FUCCI sensor (Fig. S2a) and SMAD reporter (Fig. S2b), video with in total about 2500 labelled nuclei. This dataset was recorded with a high magnification (100x) and has high signal-to-noise and signal-to-background ratios.

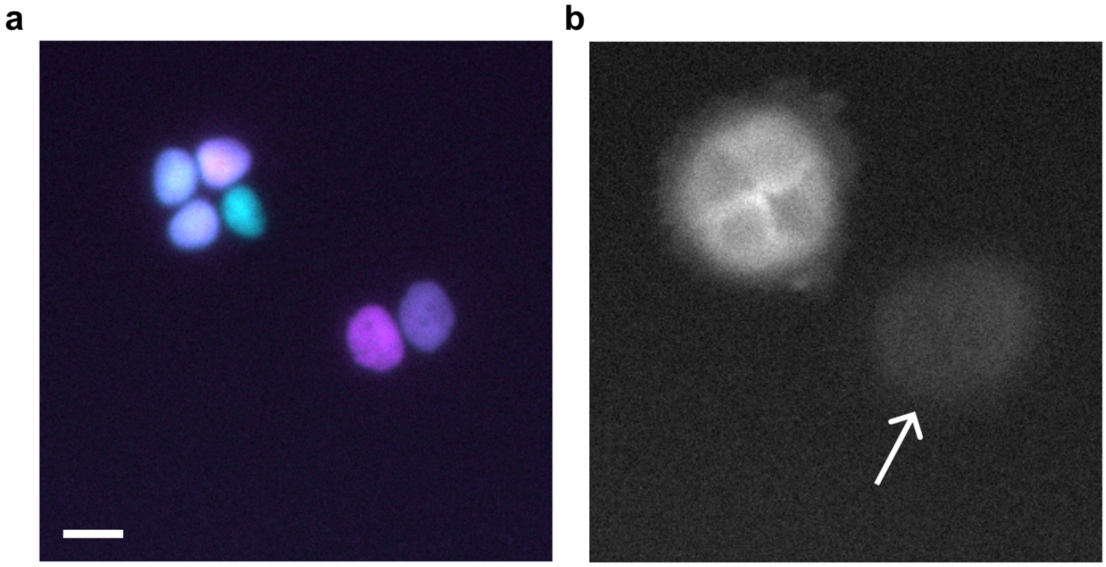


Figure S2: Example from the dataset of Han et al.[1]: **a** FUCCI signal, **b** SMAD signal. The area with low SMAD signal, where the nucleus is not clearly visible is indicated by an arrow. The scale bar is 10 µm.

### ConfluentFUCCI

Example dataset from their GitHub repository (https://github.com/leogolds/ConfluentFUCCI), we used 15 frames out of 60 with about 1000 labelled nuclei.

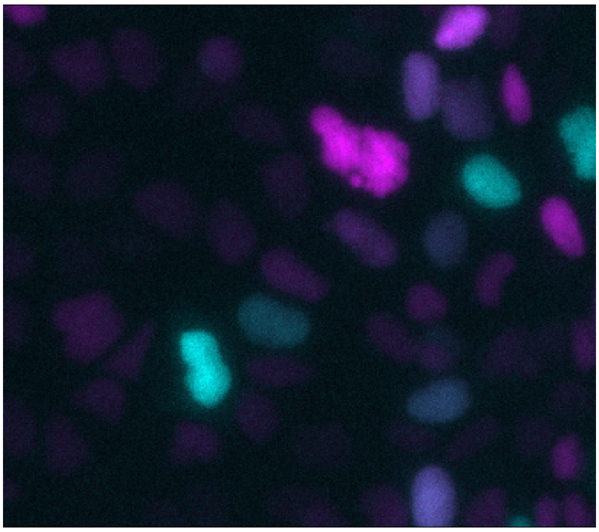

Figure S3: Example frame from the ConfluentFUCCI dataset[2]. No scale bar is provided because metadata on pixel size was missing.

#### CellMAPtracer

The dataset comprises RPE1-hTert cells labelled with PIP-FUCCI sensor. We labelled one frame containing 275 nuclei. Some of them are not visible (Fig. S4).

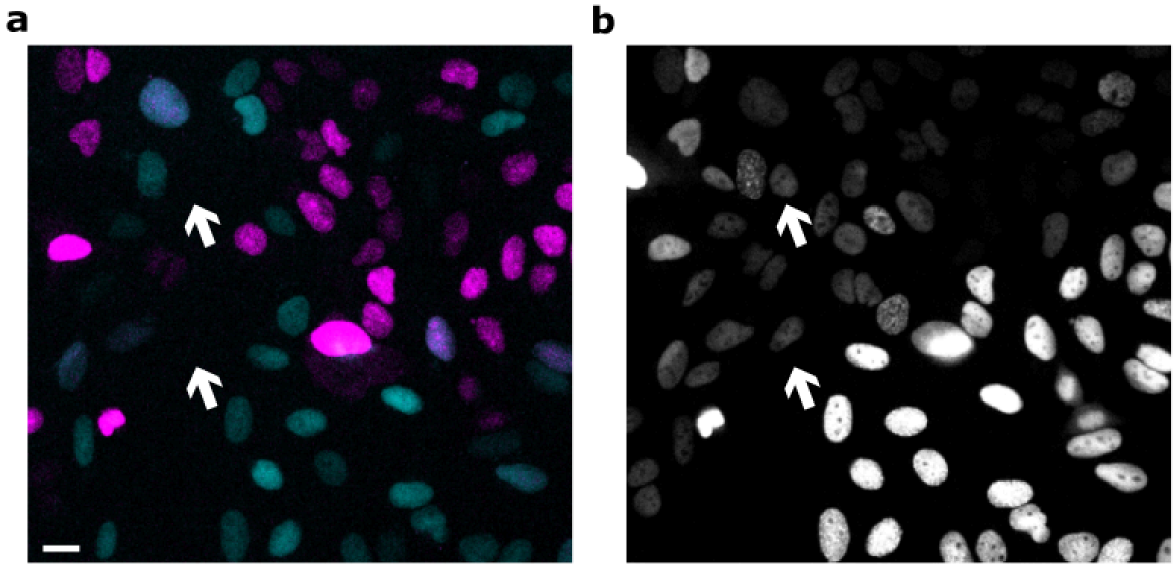


Figure S4: **a** FUCCI signal of the CellMAPtracer data[3]. **B** The nuclei were additionally labelled with PCNA because the PIP-FUCCI signal vanishes between G1 and S phase (see arrows). These nuclei were still included in the dataset and explain the in comparison lower accuracy. The scale bar is 20 µm.

### Cotton et al.

Cotton et al. presented a new PIP-H2A reporter[4]. We used one labelled frame with about 1900 labelled nuclei (see an example in Fig. S5).

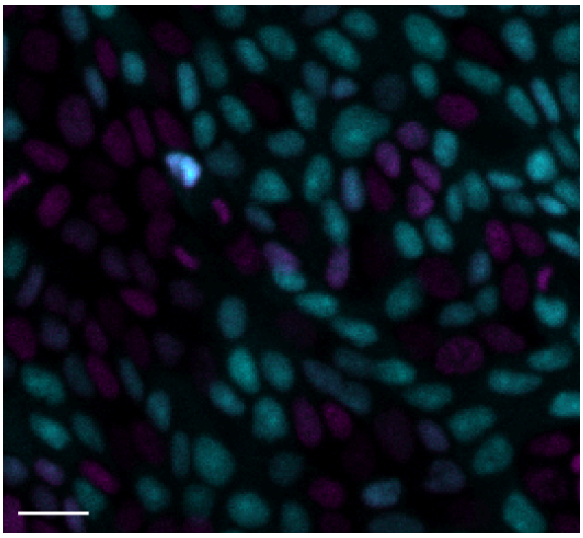

Figure S5: Example FUCCI data from Cotton et al.[4] The scale bar is 20 µm.

### Classification with Cellpose-SAM

Precision

| | Tubulin-only | | | Tubulin-only 1-CH | | | Regular FUCCI | | | Swapped FUCCI | | | All channels | | |
|---|---|---|---|---|---|---|---|---|---|---|---|---|---|---|---|
| GT \ Prediction | G1 | G1/S | S/G2/M | G1 | G1/S | S/G2/M | G1 | G1/S | S/G2/M | G1 | G1/S | S/G2/M | G1 | G1/S | S/G2/M |
| G1 | **0.25** | 0.0 | 0.0 | **0.69** | 0.29 | 0.12 | **0.89** | 0.06 | 0.00 | 0.06 | 0.18 | **0.94** | **0.90** | 0.05 | 0.00 |
| G1/S | 0.08 | 0.0 | 0.0 | 0.11 | **0.61** | 0.06 | 0.03 | **0.82** | 0.04 | 0.05 | **0.64** | 0.03 | 0.04 | **0.89** | 0.04 |
| S/G2/M | 0.35 | 0.00 | **1.0** | 0.06 | 0.07 | **0.79** | 0.01 | 0.1 | **0.96** | **0.85** | 0.17 | 0.00 | 0.01 | 0.05 | **0.94** |

Accuracy

| | Tubulin-only | | | Tubulin-only 1-CH | | | Regular FUCCI | | | Swapped FUCCI | | | All channels | | |
|---|---|---|---|---|---|---|---|---|---|---|---|---|---|---|---|
| GT \ Prediction | G1 | G1/S | S/G2/M | G1 | G1/S | S/G2/M | G1 | G1/S | S/G2/M | G1 | G1/S | S/G2/M | G1 | G1/S | S/G2/M |
| G1 | **0.18** | 0.00 | 0.02 | **0.50** | 0.02 | 0.05 | **0.82** | 0.01 | 0.00 | 0.02 | 0.04 | **0.77** | **0.83** | 0.00 | 0.00 |
| G1/S | 0.08 | **0.0** | 0.0 | 0.10 | **0.19** | 0.05 | 0.02 | **0.63** | 0.03 | 0.04 | **0.50** | 0.02 | 0.04 | **0.60** | 0.03 |
| S/G2/M | 0.32 | 0.00 | **0.0** | 0.04 | 0.01 | **0.70** | 0.01 | 0.02 | **0.90** | **0.79** | 0.05 | 0.00 | 0.00 | 0.01 | **0.91** |

**Table S1** Classification using the Cellpose-SAM network. The network was trained using all channels, only the "Tubulin-only 1CH" configuration was trained on the single tubulin channel. The network is by construction channel-invariant and can be evaluated using an arbitrary number of channels. This Table accompanies Table 2 of the manuscript.
